# Quantitative phosphoproteomic analysis of testes from *Iqcn*-deficient mice highlights the significance of calmodulin signaling in spermiogenesis

**DOI:** 10.1101/2025.03.10.642497

**Authors:** Jing Dai, Li Lou, Xingyao Wang, Yilian Huang, Jiao Lei, Feitai Tang, Yangyang Bian, Yong Zeng, Guangxiu Lu, Ge Lin, Shen Zhang

**Author notes:** Contact details for the corresponding authors: Shen Zhang, Address: 567 Tongzipo West Road, Clinical Research Center for Reproduction and Genetics in Hunan Province, Reproductive and Genetic Hospital of CITIC-XIANGYA, Changsha, Hunan 410000, China, Jing Dai, Address: 567 Tongzipo West Road, Clinical Research Center for Reproduction and Genetics in Hunan Province, Reproductive and Genetic Hospital of CITIC-XIANGYA, Changsha, Hunan 410000, China.

## Abstract

Calmodulin (CaM) plays a crucial role in sperm function. Studies have reported that proteins containing IQ motif interact with CaM, subsequently engaging with downstream target proteins known as calmodulin-binding proteins (CaMBPs). It is reported that the loss of interaction between IQCN and CaM is mainly manifested as decreased motility, leading to fertilization failure and male infertility. However, no relevant reports have been published detailing which CaMBPs exist and the mechanisms by which they are regulated. In this study, we conducted a comprehensive quantitative proteomic and phosphoproteomic analysis of mouse testes from wild-type (WT) and *Iqcn* knockout (*Iqcn*^*-/-*^) mice. The results indicated that *Iqcn* deficiency substantially rewires the downstream phosphorylation signaling pathway, while not causing equivalent changes in protein levels. Among the total 577 differentially regulated phosphorylated sites in our results, most of them (494/577) belong to calmodulin-binding proteins (CaMBPs). Gene ontology analysis of these differentially phosphorylated CaMBPs showed enrichment in male gamete generation, actin cytoskeleton organization and microtubule cytoskeleton organization process, demonstrating IQCN regulates sperm function by interacting with CaM, which in turn affects the phosphorylation level of CaMBPs. Further kinase substrate network analysis revealed that most kinases with substrates’ phosphorylation sites up-regulated were tyrosine kinases, and the inhibition assay showed that FGFR4 and SYK tyrosine kinases are important for sperm motility and progressive motility. In summary, this study reveals the interaction between IQ motif-containing protein IQCN and CaM, which regulates the phosphorylation of downstream CaMBPs and is involved in the related processes of spermiogenesis and sperm function.

## Introduction

Calmodulin (CaM) participates in various intracellular signal transduction pathways and plays a pivotal role in Ca^2+^-dependent signal transduction pathways, serving as a dynamic Ca^2+^ sensor that relays signal downstream[1]. The Ca^2+^-CaM complex interacts with target proteins and modulates their activity, a process accomplished through its interaction with downstream target proteins known as calmodulin-binding proteins (CaMBPs)[2]. CaM plays a significant role in sperm function. In mature sperms, CaM in conjunction with associated CaMBPs, plays a crucial part in processes such as sperm motility, capacitation, acrosome reaction, and sperm-egg fusion[3, 4].

Studies have shown that proteins containing the IQ motif interact with CaM, causing conformational changes in CaM and enabling it to function as a Ca^2+^ sensor[5]. The IQ motif containing genes *IQCN, IQCF1, IQCG, IQCH* and *IQCD* are functionally related to sperm function and are highly or specifically expressed in testicular tissue. Among them, IQCG is located on the manchette structure and interacts with CaM[6, 7]. *Iqcf1* is associated with sperm capacitation and acrosome reaction. The main reason for the decreased fertility of male mice with *Iqcf1* knockout is the downregulation of CaM, which affects the phosphorylation level of sperm tyrosine phosphatase, leading to decreased sperm motility and acrosome reaction rates[8]. The IQCH association with calmodulin is distributed around the sperm axoneme, serving to support its function within flagella[9]. Our previous research revealed that the knockout of *Iqcn*, a novel member of the IQ motif containing genes, caused multiple abnormalities during spermiogenesis in male mice including acrosome biogenesis and flagellum assembly, ultimately resulting in decreased sperm motility, fertilization failure and male infertility[10, 11]. However, the regulatory mechanisms of CaM on related phenotypes remain largely unclear.

For a long time, it has been speculated that protein phosphorylation and dephosphorylation play a significant role in sperm function[12]. This phenomenon is evidenced by the involvement of Calcium/calmodulin dependent protein kinase II (CAMK2) and Calcium/calmodulin dependent protein kinase IV (CAMK4) in sperm motility and hyperactivation[13]. The role of Ca^2+^/CaM is mediated through the action of kinase and phosphatases. Based on this, we hypothesize that downstream CaMBPs of CaM regulate spermiogenesis and sperm function through the process of phosphorylation and dephosphorylation. Previous studies have investigated the regulatory process of sperm maturation in the epididymis through proteomics and phosphoproteomics[14]. However, the mechanism by which the CaM pathway regulates the phosphorylation of downstream CaMBPs remains unclear.

This study employs *Iqcn* knockout mice as a model to elucidate the phosphorylation regulatory mechanism of the CaM pathway during spermiogenesis. It provides a theoretical basis for addressing the abnormalities in acrosome biogenesis, flagellum assembly, and related sperm functions caused by dysregulation of the CaM pathway.

## Experimental Section

### Samples collection

*Iqcn* knockout (*Iqcn*^*-/-*^) mice were created utilizing CRISPR/Cas9-mediated genome editing, as detailed previously[10]. In brief, guide RNAs (gRNAs) specifically targeting exon 3 and exon 4 were employed to generate *Iqcn*^*-/-*^. These gRNAs, along with Cas9 mRNA, were injected into the zygotes of C57B6J mice. The genotypes of the resulting offspring were confirmed through Sanger sequencing. Heterozygous *Iqcn*^*-/-*^ founder mice were then crossed to produce *Iqcn*^*+/-*^ mice for use in the experimental procedures.

We collected four pairs of 8 to 10-week-old wild-type and *Iqcn*^*-/-*^ male mice for proteomic and phosphoproteomic analyses. This study adhered to the recommendations outlined in the Guide for the Care and Use of Laboratory Animals and received approval from the Animal Welfare Ethics Committee of Central South University (Approval Number: XMSB-2022-0057).

### Proteomics sample preparation

Protein extraction was performed utilizing the protein extraction kit (Cat # SA-04-TD, Invent) according to the manufacturer’s protocol. Both the protease inhibitor cocktail (Cat # 78429, Thermo Fisher Scientific) and phosphatase inhibitors (Cat # 4906845001, Roche) were added into the lysis buffer. The protein concentration was then quantified using BCA assay kit (Cat # 23225, Thermo Fisher Scientific)

Equal amount of proteins for each sample were transferred to centrifugal filters (Cat # VN01H02, Sartorius) with 10K molecular weight cut off and washed three times with 6 M guanidine hydrochloride in 25□mM Triethylammonium bicarbonate (TEAB). Then the protein samples were reduced by 100 mM DL-dithiothreital (DTT) (Catalog # D9779, Sigma-Aldrich) for 1 h at 56 °C, followed by alkylation of thiol group with 100 mM Iodoacetamide (IAA) (Cat # I1149, Sigma-Aldrich) for 40min in the darkness at room temperature. Afterward, background solution was exchanged to 25□mM TEAB by centrifugation and the sample solution was subjected to proteolytic digestion with sequencing-grade trypsin (Cat # HLS TRY001C, Hualishi Scientific) at 1:25 (m/m) enzyme-to-substrate ratio for 16 hours at 37 °C. After digestion, the filters were centrifuged at 14,000 g for 15 min, and the flow-through containing the peptides was collected. To increase peptide recovery from the membrane, the membrane was further washed with water and the flow-through from those two steps was then combined. At last, the flow-through was lyophilized for subsequent proteomics analysis and phosphopeptide enrichment.

### Phosphopeptide Enrichment

The lyophilized peptides were selectively enriched with High-Select™ TiO_2_ Phosphopeptide Enrichment Kit (Cat # A32993, Thermo Fisher Scientific) according to the manufacturer’s guidelines. The enriched phosphopeptides were lyophilized for LC-MS/MS analysis.

### LC-MS/MS Analysis

The MS analysis for digested peptides and enriched phosphopeptides were both performed according to our recent study[15]. Briefly, separation was performed on a 25 cm PepMap analytical column (ThermoFisher Scientific) with 3% to 38% buffer B (80% ACN, 0.1% formic acid) for 102 min and kept at 100% buffer B for 10min. FAIMS switched between CVs of −35 V, −45V and −60 V with 1 s cycle time. MS1 spectra were obtained in the Orbitrap (resolution: 60k; AGQ target: standard; MaxIT: Auto; RF lens: 50%; mass range: 350 to 1,500). Dynamic exclusion was used for 20 s to exclude all charge states for a specified precursor. The collection of MS2 spectra was completed in the linear ion trap (isolation window: 1.6 m/z; scan rate: rapid; AGQ target: standard; MaxIT: Auto; HCD CE: 35%; data type: centroid).

### Database searching

Proteome Discoverer Software (version 2.5, San Jose, CA) was used to process raw files for detecting features, searching databases and quantifying proteins/peptides. MS/MS spectra were searched against the UniProt mouse database (downloaded on June 24th, 2022, containing 55,286 entries). Methionine oxidation and N-terminal protein acetylation were chosen as variable modifications, while the carbamidomethylation of cysteine residues was regarded as a fixed modification. Precursors and fragments had a mass tolerance of 10 ppm and 0.6 Da, respectively. Minimum and maximum peptide lengths were six and 144 amino acids, respectively. The missed cleavage allowed for every peptide was two. The filtering of proteins had a maximum false discovery rate (FDR) of 0.01. The default settings of Proteome Discoverer were used for other parameters that were not mentioned.

The phosphoproteomics data was analyzed using MaxQuant software (version 2.1.0.0) and searched against the UniProt mouse database (downloaded on June 24th, 2022, containing 55,286 entries). Trypsin was selected as the enzymatic digestion, and a maximum of two missed cleavages was allowed. The mass tolerance for first search and main search were set to 20 ppm and 4.5 ppm. Carbamidomethylation (Cys) was set as the fixed modification. Variable modifications were set to oxidation (M), acetylation (protein N-terminus) and phosphorylation (STY). False discovery rate (FDR) cut-offs were set to 0.01 for proteins, peptides, and phosphorylation sites to control the rate of false identification. Only phosphorylation sites with localization probability larger than 0.75 were retained in downstream analysis.

### Data analysis

The phosphoproteomics data acquired by label-free quantification (LFQ) were analyzed as follows: firstly, phosphosites were filtered based on their quantification across mice requiring at least 50% of quantification in at least one condition. Since the data were acquired as LFQ, a stepwise approach implemented in PhosR [16] was applied to impute missing values, including site- and condition-specific imputation (scImpute) and paired-tail imputation (ptImpute). Differentially regulated phosphosites from normalized data between wild-type and *Iqcn*^*-/-*^ groups were filtered by p < 0.05 & fold change > 1.5 with Limma R package [17] using paired analysis. Similar with phosphoprteomics data, proteomic data from wild-type and *Iqcn*^*-/-*^ groups were log_2_ transformed, median centered and scaled, filtered and subjected to Limma differential expression analysis using the same thresholds.

Gene ontology (GO) and Kyoto Encyclopedia of Genes and Genomes (KEGG) pathway enrichment analysis were performed utilizing Metascape (v3.5) [18] with *P* value less than 0.05 as a cut-off. Phenotypes of the differentially phosphorylated proteins were annotated according to the MGI database (v6.24) with *P* value less than 0.05 [19]. Kinase enrichment analysis was executed using PhosphoSitePlus (v6.7.5) [20], contrasting significantly up-regulated or down-regulated phosphorylation sites against the remainder of phosphorylation sites. Kinome tree was generated using KinMap [21].

### Immunoblotting

Protein lysates were boiled at 100°C for 10 minutes in SDS-PAGE loading buffer (FDB10), subsequently separated by 10% SDS-PAGE, and transferred onto polyvinylidene difluoride (PVDF) membranes. The membranes were blocked with 5% BSA (bovine serum albumin) in TBST and then incubated overnight at 4 ° C with primary antibodies, specifically Anti-Phospho-Tyrosine (PTM BIO, PTM-702RM) and Anti-GAPDH (ABclonal, AC033). Following this, the membranes were washed three times with TBST and incubated with secondary antibodies, Goat Anti-Rabbit IgG H&L (HRP) from Abcam, for 2 hours at room temperature. Protein bands were visualized using the High-sig ECL Western Blotting Substrate (Beyotime, P0018FS).

### Kinase inhibition assays

Mouse sperm were released from the epididymides into 1 ml of human tubal fluid (HTF; Aibei Biotechnology, M1130, Nanjing, China) medium for 15min at 37°C. Following this, the sperm suspension from wild-type male mice was evenly divided into three replicate portions. Each portion was then supplemented with a final concentration of 3nM BLU9931 (FGFR4 inhibitor, Selleck, S7819, China), 41 nM R406 (SYK inhibitor, Selleck, S2194, China), or 0.02% dimethyl sulfoxide (DMSO) as a control. The sperm suspension from *Iqcn*^*-/-*^ male mice was supplemented with 0.02% DMSO. Subsequently, mouse sperm motility was evaluated using a computer-assisted semen analysis system (SAS medical, SAS-MS). The sperm motility parameters VCL (curvilinear velocity), VSL (straight-line velocity) and VAP (average path velocity) were examined.

## Results

### Quantitative proteomics and phosphoproteomic analysis of testes from *Iqcn*-deficient mice

To investigate the regulatory mechanism of IQCN and its signaling pathway during spermiogenesis, quantitative proteomics and phosphoproteomic analysis of testes from *Iqcn*^*-/-*^ mice were performed in four replicates (Figure 1A). In the quantitative proteomics results, a total of 6,264 proteins were identified in our mouse testes samples, among them only 86 proteins were up-regulated and 72 proteins were down-regulated in *Iqcn*^*-/-*^ mice (supplemental Table S1), revealing the expression of most proteins were not significantly changed with *Iqcn* deficiency. For quantitative phosphoproteomics analysis, a total of 8,500 phosphorylation sites corresponding to 5,039 phosphopeptides and 2,963 phosphoproteins were identified in our results (supplemental Table S2). Among the identified phosphorylation sites, 6,235 were class I phosphorylation sites with localization probability above 0.75, corresponding to 4,496 phosphopeptides and 2,756 phosphoproteins (supplemental Table S3). The 6,235 Class I sites comprised 79.6% phosphoserine (pS), 11.3% phosphothreonine (pT), and 9.1% phosphotyrosine (pY) (Figure 1B). It is worth to note that, although the pY ratio in our study is much higher than that in the reported works (Figure S1A)[22, 23], most of them contain the diagnostic peak (Figure S1B), suggesting the reliability of the results and indicating the *Iqcn* knockout may cause alteration in tyrosine phosphorylation. Compared with the PhosphositePlus database (https://www.phosphosite.org), we identified 3406 new phosphorylation sites that have not been reported before, accounts for 54.6% of all phosphorylation sites identified by this study (Figure 1C). Comparative analysis of our proteomes and phosphoproteomes provides the opportunity to explore whether altered downstream signaling coincided with changes in protein abundance. Based on the quantification results on protein level and phosphorylation sites level, we performed the principal component analysis (PCA), hierarchical clustering and pearson’s correlation analysis (Figure 1D-I). These results firstly further suggested that *Iqcn* deficiency caused limited perturbation on protein expression level as WT and *Iqcn*^*-/-*^ proteome can’t be divided into different groups from PCA and hierarchical clustering, and the correlation coefficients between different groups were generally similar. Secondly, these results indicated different phosphorylation state between these two groups, reflected by the distinct clusters from hierarchical clustering and higher correlation coefficients within groups compared with those between groups. The above conclusion was further confirmed by differential expression analysis of proteome and phsophoproteome, as well as the correlation analysis between them (Figure 1J-L), in which only 2.52% (158/6,264) of the total identified proteins were differential regulated while more than 9.25% (577/6,235) of the phosphorylation sites showed significant changed, and poor correlation between changes in phosphoproteome and proteome expression. Overall, our findings imply that *Iqcn* deficiency substantially rewires the downstream signaling pathway, and this is generally not due to equivalent changes in protein levels.

**Figure 1.**
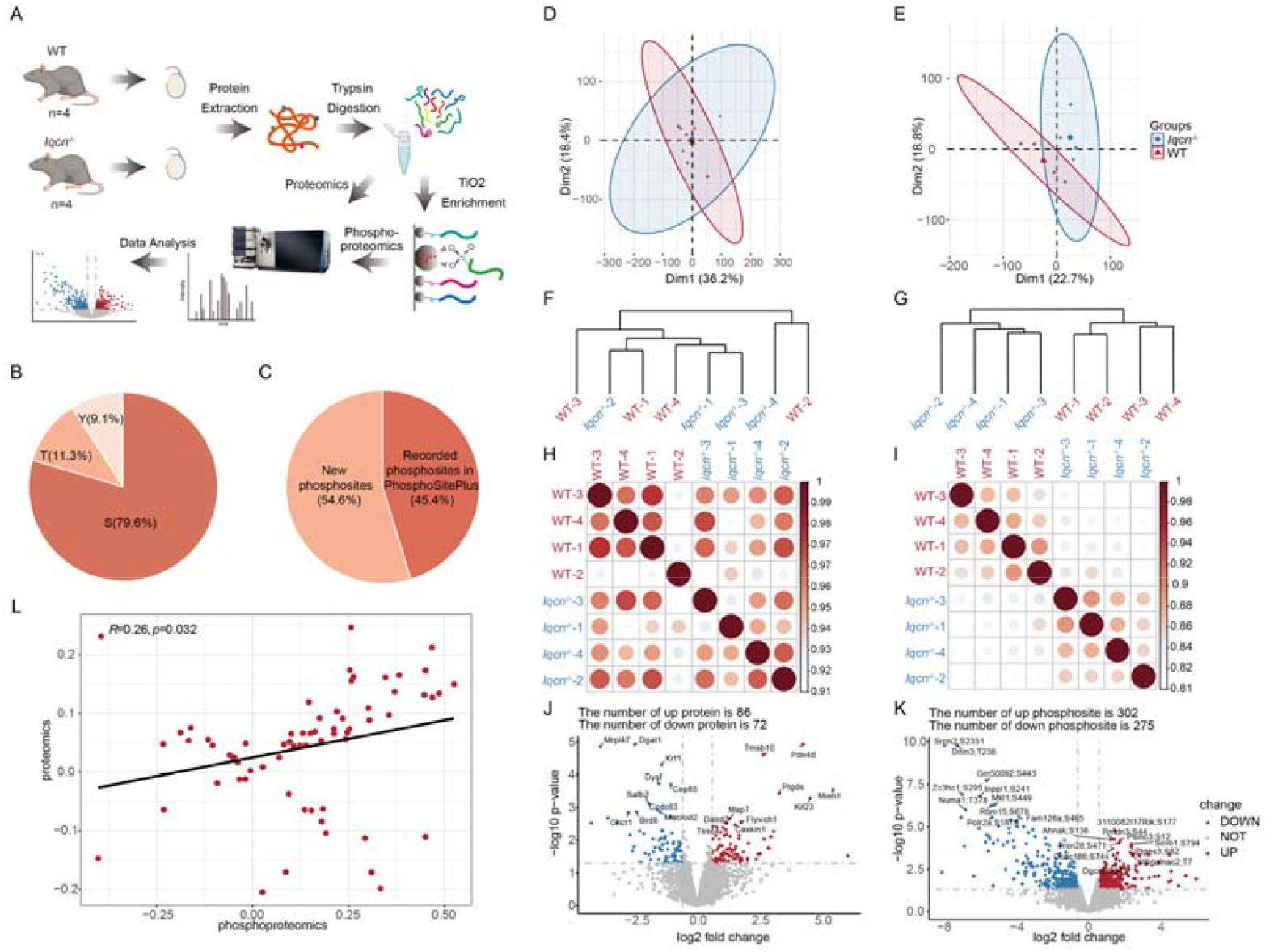
Quantitative proteomics and phosphoproteomic analysis of testes from *Iqcn*-deficient mice. (A) Proteomics and phosphoproteomic analysis workflow for testes from wild-type (WT) and *Iqcn* knockout mice. (B) Distribution of phosphorylation on serine (S), threonine (T), and tyrosine (Y) for identified phosphopeptides. (C) Comparison of the phosphosites identified here with those recorded in PhosphositePlus. (D) Principal component analysis (PCA) based on proteomics data. (E) PCA based on phosphoproteomics data. (F) Hierarchical clustering based on proteomics data. (G) Hierarchical clustering based on phosphoproteomics data. (H) Pearson’s correlation between different proteomics analysis replicates. (I) Pearson’s correlation between different phosphoproteomics analysis replicates. (J) Differential regulated proteins between testes from WT and *Iqcn* knockout mice. (K) Differential regulated phosphosites between testes from WT and *Iqcn* knockout mice. Red: up-regulated, blue: down-regulated, fold change ≥1.5, p ≤ 0.05. (L) Correlation of proteome and phosphoproteome.

### Functional analysis of differentially regulated phosphoproteins during spermiogenesis

To investigate the regulatory mechanism of the CaM on downstream protein and signaling pathway during spermiogenesis, we compared our differential phosphoproteome data with Calmodulin Target Database (http://calcium.uhnres.utoronto.ca/ctdb/ctdb/home.html) and found 380 calmodulin-binding proteins (CaMBPs) in our dataset, corresponding to 494 differential regulated phosphorylation sites, which accounted for 85.62% (494/577) of all differential regulated phosphorylation sites in our phosphoproteomics result, including 256 up-regulated sites and 238 down-regulated sites (supplemental Table S4). Among them, 7 proteins including FSIP2, ADGB, REEP6, AKAP3, HGSNAT, PPP1R7 and SRRM2 had both up- and down-regulated phosphorylation sites, suggesting that these proteins undergo complex phosphorylation regulation at different locations in the testes (Figure S2). For example, FSIP2 has one downregulated phosphorylation site (S6986) and one upregulated phosphorylation site (T592). Previous studies have demonstrated that alterations in the phosphorylation of FSIP2 in sperm are associated with cryopreservation-induced decreases in sperm motility[24], thereby providing high confidence in our quantitative phosphoproteomics data.

To characterize the functions of these differentially phosphorylated CaMBPs, we performed gene ontology enrichment analysis (Figure 2A and supplemental Table S5). We found that enrichment of terms related to male gamete generation, actin cytoskeleton organization and microtubule cytoskeleton organization process (Figure 2A). These proteins were involved in sperm flagellum formation, such as up-regulated proteins SLC26A8, MAP1B, KIF9, PCM1, ADGB and ENKD1 and down-regulated proteins ALS2CR11, DYNC1I2, SPATA18, CFAP45, SPAG17, DNAH7A, PCM1, ADGB and CATSPERG2 (Figure 2B). It also involved in sperm acrosome formation, such as AKAP3, FSIP2, CSNK2A2, PDCL2, RIMBP3 and VAPS13B (Figure 2C). Then, we also observed enrichment of terms in sperm capacitation and fertilization. For example, it contained YBX2, IZUMO1 and SLC4A2 (Figure 2D). These results indicated changes in the phosphorylation levels of a series of CaMBPs might be involved in a range of biological events, including acrosome formation, flagellum development, and the fertilization process in spermatozoa.

**Figure 2.**
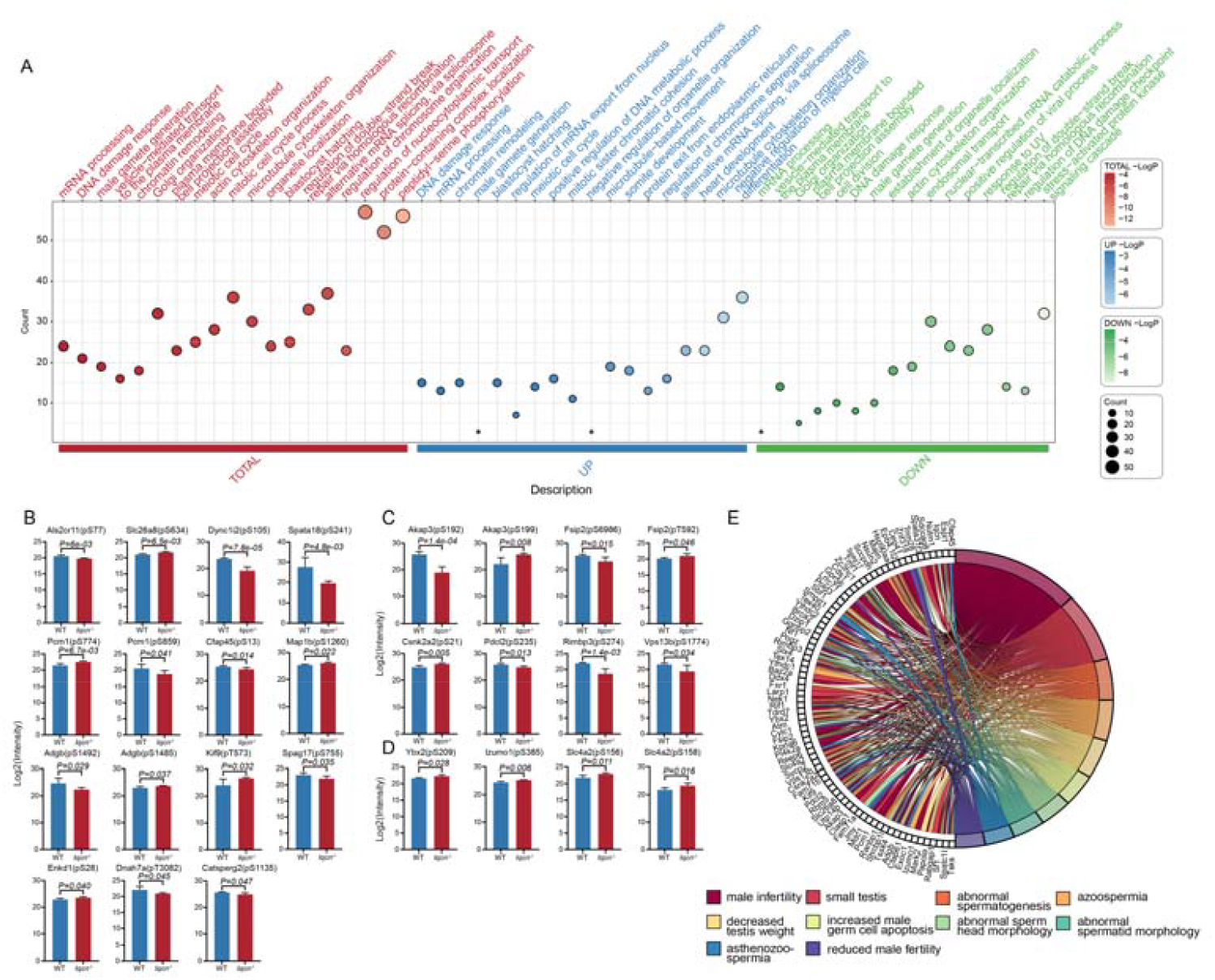
Functional analysis of differentially regulated phosphoproteins during spermiogenesis. (A) Gene ontology enrichment analysis of differentially phosphorylated proteins. (B) The quantification of differentially regulated phosphorylation sites of sperm flagellum related proteins. (C) The quantification of differentially regulated phosphorylation sites of acrosome related proteins. (D) The quantification of differentially regulated phosphorylation sites of fertilization related proteins. (E) MGI male reproductive phenotypes related differentially phosphorylated proteins.

To further analyze the *in vivo* functions of proteins in mouse, we annotated phenotypes of the differentially phosphorylated proteins according to the MGI database (https://www.informatics.jax.org/) (Figure 2E). The results showed that proteins with altered levels of phosphorylation during the spermatogenesis and spermiogenesis were closely related to male infertility and sperm abnormalities. In total, 56 proteins were annotated to male infertility, 19 proteins were annotated to abnormal spermatogenesis, 19 proteins were annotated to azoospermia, 14 proteins were annotated to abnormal sperm head morphology and 13 proteins were annotated to asthenozoospermia (Figure 2E). The phosphorylation changes of these proteins which are essential for male fertility indicated the potential importance of phosphorylation regulations in sperm morphology and sperm function.

### Kinase analysis of sperm phosphoproteome during spermiogenesis

Protein phosphorylation is a reversible dynamic process that is regulated by the competing activities of protein kinases and phosphatases. With the quantitative testes proteome and phosphoproteome data, we analyzed the expression and phosphorylation levels of kinases and phosphatases in testes. In our quantitative proteomic dataset, we found that among the quantified 255 kinases and 138 phosphatases in testes, only 7 kinases and 2 phosphatases showed differential expression (Figure 3A, supplemental Table S1).

**Figure 3.**
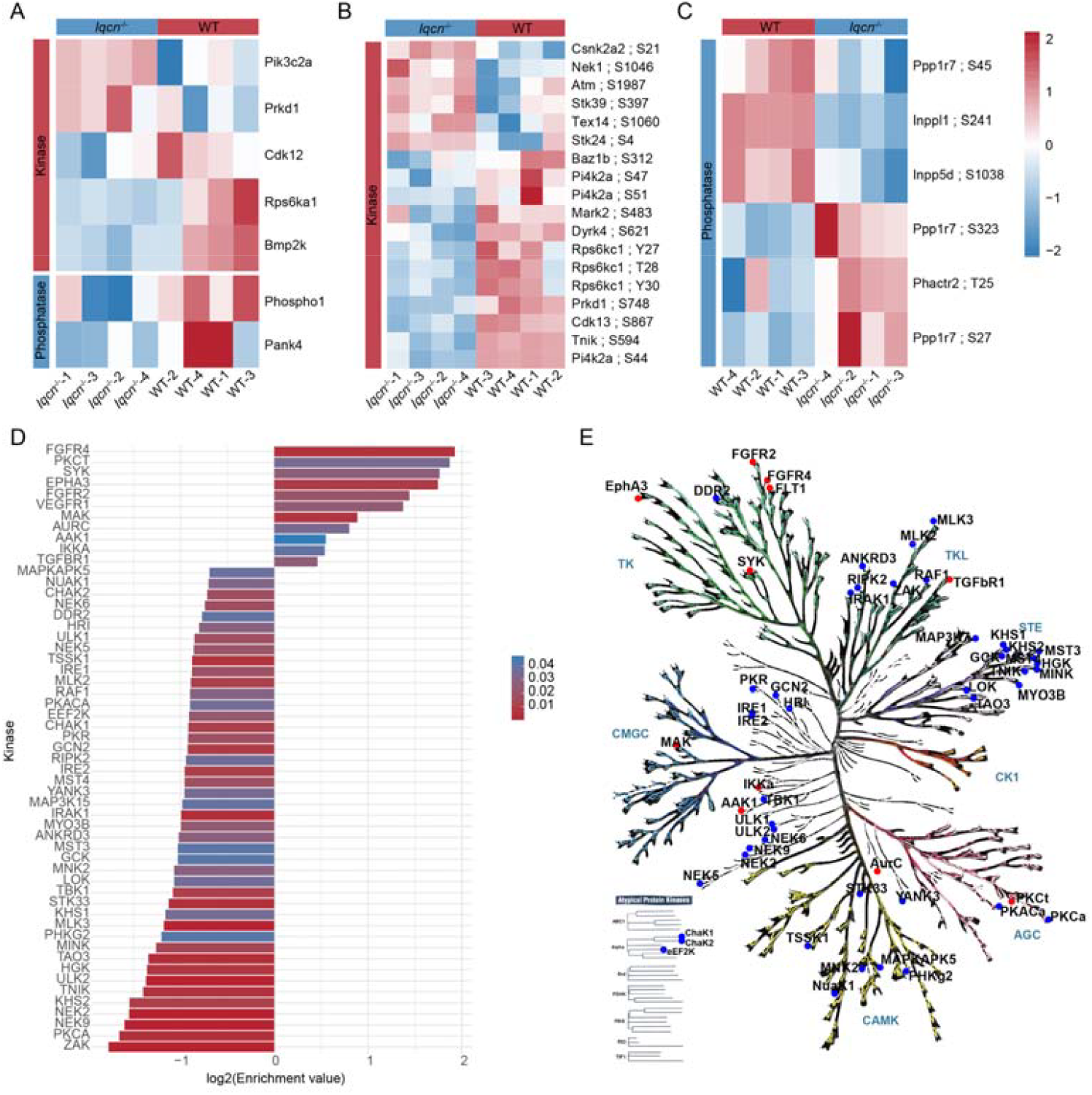
Kinase analysis of testes phosphoproteome during spermiogenesis. (A) Differentially regulated expression of kinases and phosphatases in testes during spermiogenesis. (B) Differentially regulated phosphorylation sites of kinases and phosphatases during spermiogenesis. (C) The top 20 kinases with enriched phosphorylation substrates in testes (P<0.05). (D) Enrichment of kinases with upregulated phosphorylation sites (red) and kinases with downregulated sites (blue) during spermiogenesis.

Kinases and phosphatases may also be regulated by phosphorylation, we therefore analyzed phosphorylation changes of kinases and phosphatases in our quantitative phosphoproteomic dataset. The results showed that among 319 phosphorylation sites from 149 kinases and 78 phosphorylation sites from 41 phosphatases, 18 sites from 15 kinases and 6 sites from 4 phosphatases in the 380 CaMBPs showed significant changes (Figures 3B and 3C, supplemental Table S4).

Kinases often recognize certain motifs to phosphorylate their specific substrate proteins. With the sequences surrounding phosphorylation sites, it is possible to annotate the kinase-substrate phosphorylation relationship. Utilizing PhosphoSitePlus, we predicted kinase-substrate relationships and identified kinases with enriched phosphorylation substrates (Figure 3D and supplemental Table S6). Our findings indicated that a majority of kinases exhibiting up-regulation in phosphorylation sites were tyrosine kinases. whereas most kinases with enriched phosphorylation sites down-regulated belonged to the CMGC, CAMK, STE and TKL families (Figure 3E).

### Inhibition of FGFR4 and SYK suppresses sperm motility

Given that kinases with multiple regulated phosphorylation substrates in kinase-substrate analyses may play vital roles in regulating sperm motility, we did a further examination of the expression levels and available inhibitors for the predicted kinases. Our results indicated that FGFR4 and SYK are present in the sperm proteome and have specific inhibitors that are appropriate for functional experiments.

We collected sperm from the cauda epididymis, and incubated the sperm with FGFR4 inhibitor BLU9931 or SYK inhibitor R406. BLU9931 is a potent and selective inhibitor of FGFR4 kinase with an IC50 of 3 nM. R406 is a potent inhibitor of SYK kinase with an IC50 of 41 nM.

Compare with sperm from *Iqcn*^*+/+*^ male mice, we found that the motility (51.67±1.1 v.s. 42.95±4.2 and 42.89±2.1, P=0.02) and progressively motility (11.13±0.70 v.s. 8.13±2.2 and 8.1±1.2, P=0.01) of sperm from *Iqcn*^*+/+*^ male mice after treatment with inhibitors R406 and BLU9931 significantly decreased, consistent with the motility ability observed in *Iqcn*^*-/-*^ mice (Figures 4A and 4B). Furthermore, by analyzing the movement trajectories of the sperms, it was found that their average path velocity (VAP, 13.8±0.9 v.s. 11.3±2.3 and 12.1±0.9, P=0.04) and straight-line velocity (VSL, 8.9±0.8 v.s. 7.3±1.7 and 7.8±0.6, P=0.04) also significantly decreased. The above results indicated that both SYK and FGFR4 play a role in the process of spermiogenesis by regulating the phosphorylation level of calmodulin-binding proteins, thereby influencing the motility of sperms.

**Figure 4.**
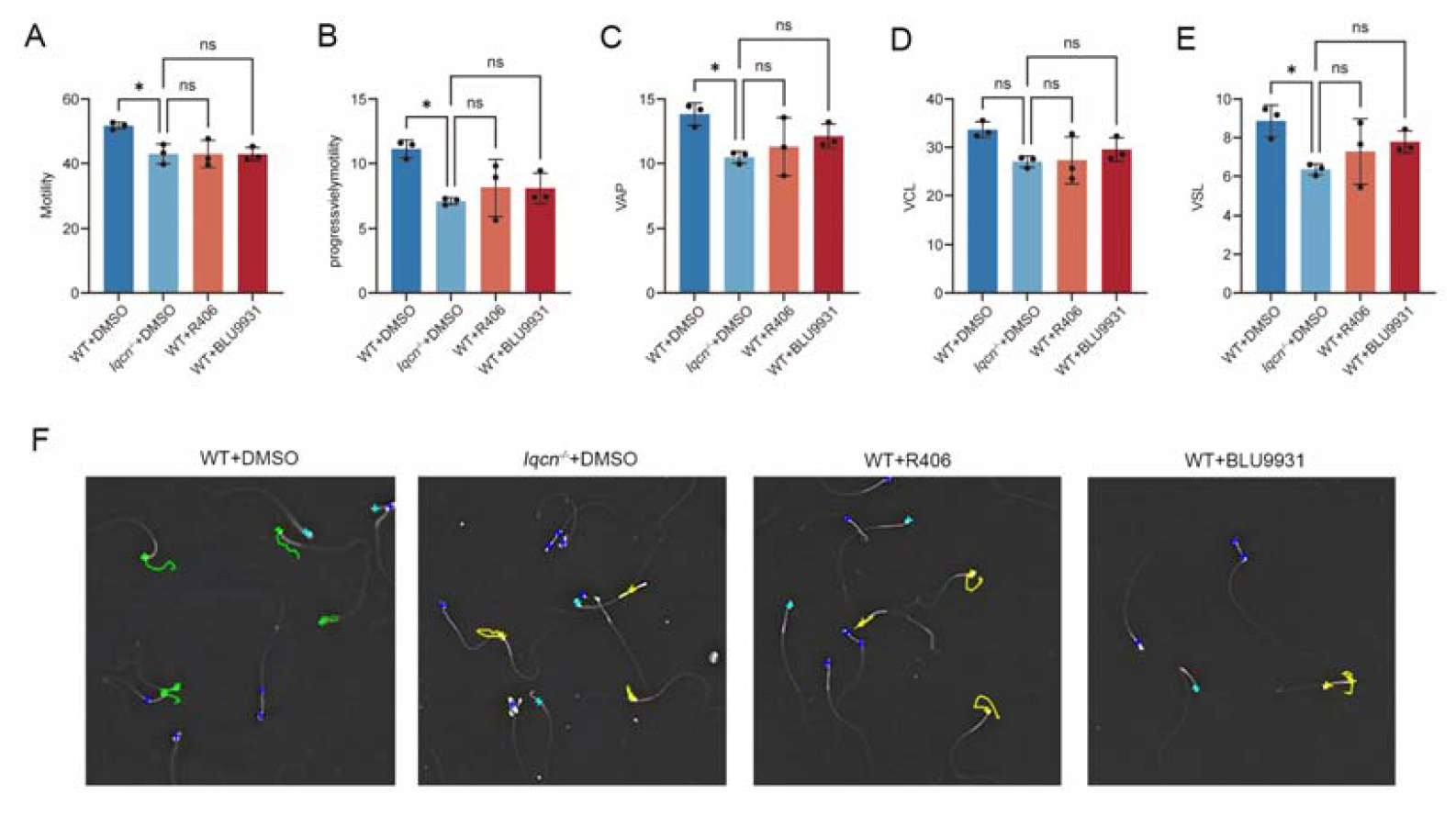
Inhibition of FGFR4 and SYK kinase affects sperm motility and progressively motility. (A-E) Sperm motility (A), progressively motility (B), VAP (C), VCL (D) and VSL (E) analyses using BLU9931 or R406 inhibitor to inhibit FGFR4 or SYK, respectively, with *Iqcn*^*+/+*^+DMSO as a positive control and *Iqcn*^*-/-*^+DMSO as a negative control. *p<0.05, n.s., not significant; one-way ANOVA. (F) The sperm motility trajectories obtained after treatment were measured using CASA. Green represents the trajectories of rapidly moving forward sperm, yellow represents the trajectories of slowly moving forward sperm, sky blue represents the trajectories of sperm moving in place, and dark blue represents the trajectories of non-moving sperm.

## Discussion

During the process of spermiogenesis, the IQ motif interacts with CaM to regulate a series of biological events, such as acrosome formation, flagella assembly, and sperm motility. However, the precise mechanism by which the IQ motif regulates downstream CaMBPs through its interaction with CaM remains largely unknown. We conducted the proteome and phosphoproteome analysis of testes from WT and *Iqcn*^*-/-*^ male mice, and identified 577 class I differentially regulated phosphorylation sites corresponding to 441 phosphoproteins. These regulated phosphoproteins are enriched in male gamete generation, actin cytoskeleton organization and microtubule cytoskeleton organization process. Kinase-substrate relationship analysis followed by functional studies confirmed the important regulatory roles of FGFR4 and SYK tyrosine kinases in sperm motility regulation. Our research demonstrates that the CaM signaling pathway is essential in the process of spermiogenesis in mice with *Iqcn* deficiency.

In our previous studies utilizing immunoprecipitation-mass spectrometry (IP-MS) for CaM in *Iqcn*^*-/-*^ male mice, we identified related CaMBPs involved in pathways such as microtubule movement, negative regulation of cytoskeleton organization, microtubule bundle formation, and actin cytoskeleton organization[10]. These proteins are highly overlapped with those identified by mass spectrometry analysis after IQCN-IP[11]. This suggests that proteins containing the IQ motif may regulate downstream CaMBPs through their interaction with CaM, thereby influencing the biological events related to spermiogenesis. However, we have observed slight phenotypic differences in male infertility caused by different IQ motif family genes. It is speculated that these differences may arise from the distinct structures of various IQ motif family members in binding to CaM, leading to different downstream CaMBP binding partners.

Using kinase-substrate prediction, we observed that most of the upregulated kinases are tyrosine kinases, prompting us to hypothesize that the IQ motif primarily modulates sperm-related functions via tyrosine kinases. It was reported that tyrosine phosphorylation and its upregulation by cAMP have been associated with capacitation and motility changes of spermatozoa[25]. We have found that inhibitors of FGFR4 and SYK kinases have the effect of inhibiting sperm motility. We discovered that these two kinases both phosphorylate and activate the PLCG1 signaling pathway. Reports indicate that zona pellucida proteins bind to tyrosine kinase receptors, leading to the phosphorylation of Ca^2+^ transporters and the entry of Ca^2+^. Also, PLCG1 is phosphorylated and upon increase of Ca^2+^ activated. Consequently, PLCG1 is translocated and coupled to the plasma membrane. This provides a theoretical basis for IQCN’s involvement in the phosphorylation of sperm tyrosinase, thereby regulating the ability of sperm to fertilize.

## Conclusions

In summary, this study conducted proteomic and phosphoproteomic analyses on testicular tissues collected from mice with *Iqcn* deficiency, revealing the process of IQ motif in regulating the CaM signaling pathway and the phosphorylation of its downstream proteins. This provides a theoretical basis for further elucidating the mechanisms related to sperm function during spermiogenesis.

## ASSOCIATED CONTENT

### Data Availability Statement

The proteomics data have been deposited in the ProteomeXchange Consortium via the proteomics identifications (PRIDE) database (Identifier: PXD057904).

### Supporting Information

The Supporting Information is available, free of charge at…

Distribution of phosphorylation on serine (S), threonine (T), and tyrosine (Y) and the ratio of tyrosine phosphorylation sites with diagnostic peak (Figure S1); Proteins with both upregulated and downregulated phosphorylation sites (FigureS2) (PDF)

Quantitative proteomics results (Table S1) (XLSX)

Phosphoproteomics results (Table S2) (XLSX)

Class I Phosphorylation sites quantified in our results (Table S3) (XLSX)

Class I Phosphorylation sites quantified in our results (Table S3) (XLSX)

Differential regulated phosphorylation sites in calmodulin-binding proteins (Table S4) (XLSX)

Gene ontology enrichment analysis of differentially phosphorylated CaMBPs (Table S5) (XLSX)

Kinase with enriched substrate phosphorylation sites differentially regulated (Table S6) (XLSX)

### Notes

The authors declare no competing financial interest.

## Ackonwledgments

The authors thank Prof. Anne-Claude Gingras for her valuable suggestions. This work was supported by the National Natural Science Foundation of China (82202053, 32273111 and 22274130), the Scientific Research Foundation of Reproductive and Genetic Hospital of CITIC-Xiangya (YNXM-202201, YNXM-202211,YNXM-202313 and YNXM-202411), the Hundred Youth Talents Program of Hunan Province and the Hunan Provincial Grant for Innovative Province Construction (2019SK4012).

