## Supplemental Figure for "Quantitative phosphoproteomic analysis of testes from *Iqcn*-deficient mice highlights the significance of calmodulin signaling in spermiogenesis"

Shen Zhang

Address: 567 Tongzipo West Road, Clinical Research Center for Reproduction and Genetics in Hunan Province, Reproductive and Genetic Hospital of CITIC-XIANGYA, Changsha, Hunan 410000, China

Jing Dai

Address: 567 Tongzipo West Road, Clinical Research Center for Reproduction and Genetics in Hunan Province, Reproductive and Genetic Hospital of CITIC-XIANGYA, Changsha, Hunan 410000, China

**Table of Contents**

1. **Figure S1. Distribution of phosphorylation on serine (S), threonine (T), and tyrosine (Y) and the ratio of tyrosine phosphorylation sites with diagnostic peak...............................................................................................................................................S3**
2. **Figure S2. Proteins with both upregulated and downregulated phosphorylation sites................................................................................................................................................S4**

**Figure S1.** (A) Distribution of phosphorylation on serine (S), threonine (T), and tyrosine (Y) in other works and in ours. (B) The ratio of tyrosine phosphorylation sites with diagnostic peak in the database searching results.

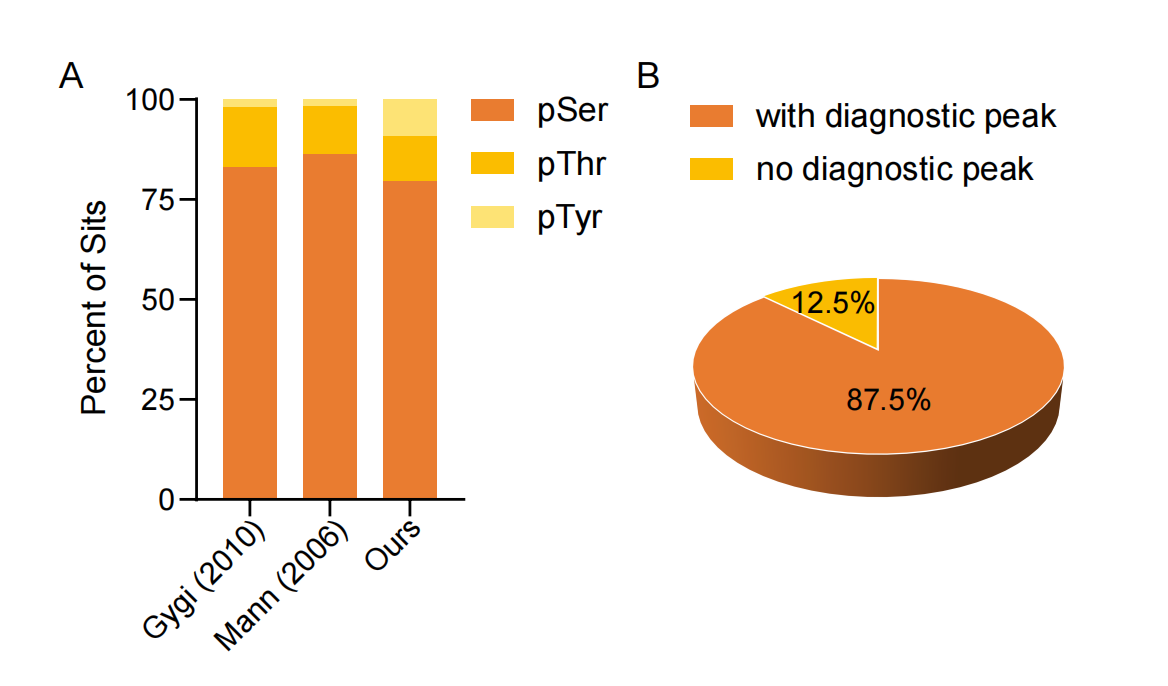

**Figure S2.** Proteins with both upregulated and downregulated phosphorylation sites.

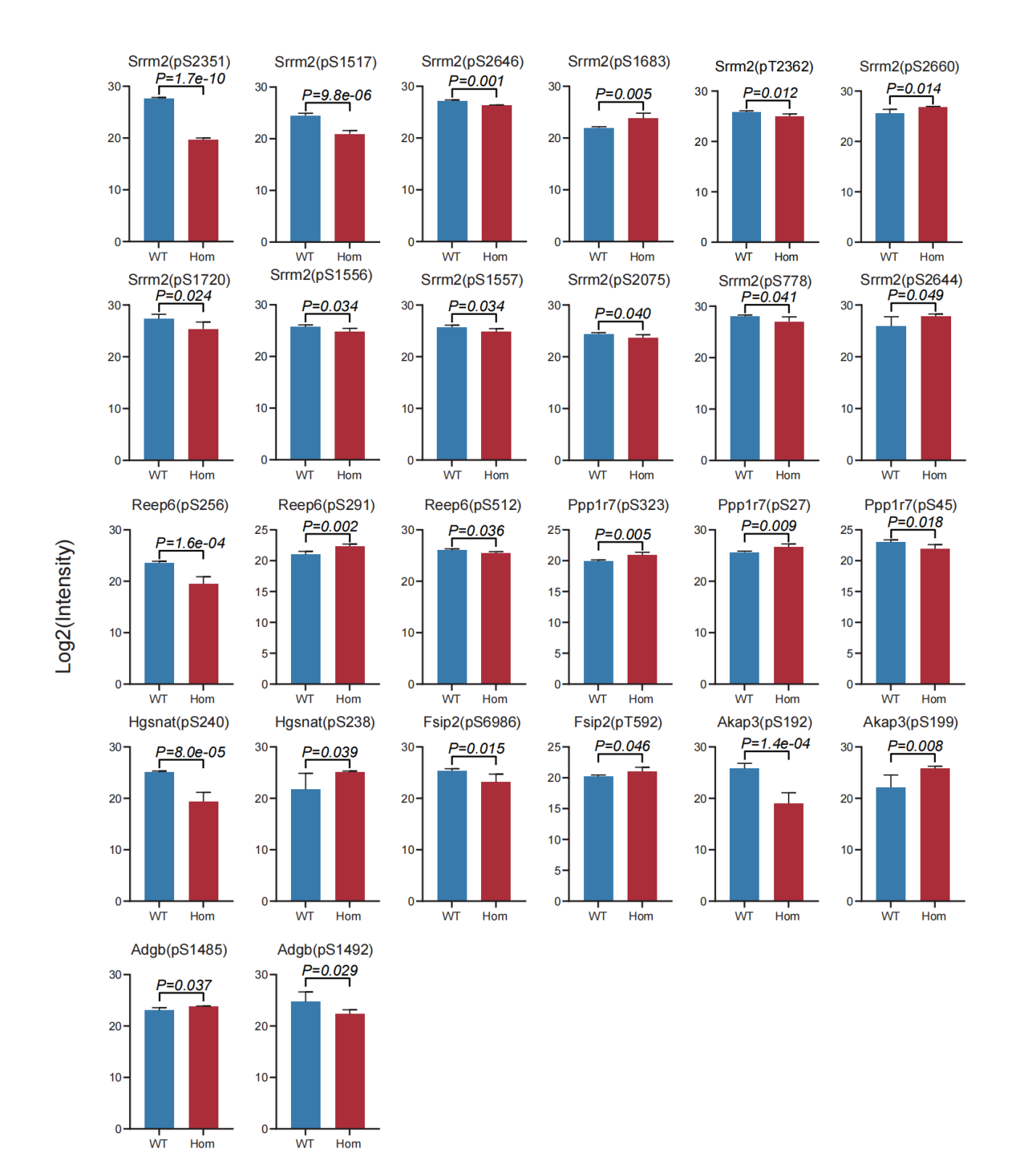
